# Searchlight-based trial-wise fMRI decoding in the presence of trial-by-trial correlations

**DOI:** 10.1101/2023.12.05.570090

**Authors:** Joram Soch

## Abstract

In multivariate pattern analysis (MVPA) for functional magnetic resonance imaging (fMRI) signals, trial-wise response amplitudes are sometimes estimated using a general linear model (GLM) with one onset regressor for each trial. When using rapid event-related designs with trials closely spaced in time, those estimates can be highly correlated due to the temporally smoothed shape of the hemodynamic response function. In previous work (Soch, J., Allefeld, C., & Haynes, J.-D. (2020). Inverse transformed encoding models – a solution to the problem of correlated trial-by-trial parameter estimates in fMRI decoding. *NeuroImage*, 209, 116449, 1-19. https://doi.org/10.1016/j.neuroimage.2019.116449), we have proposed inverse transformed encoding modelling (ITEM), a principled approach for trial-wise decoding from fMRI signals in the presence of trial-by-trial correlations. Here, we (i) perform simulation studies addressing its performance for multivariate signals and (ii) present searchlight-based ITEM analysis – which allows to predict a variable of interest from the vicinity of each voxel in the brain. We empirically validate the approach by confirming *a priori* plausible hypotheses about the well-understood visual system.

## 1 Introduction

In functional magnetic resonance imaging (fMRI), data are frequently analyzed with univariate encoding models (Brodersen et al., 2011b) such as general linear models (GLMs) as well as multivariate decoding algorithms (Brodersen et al., 2011a) such as support vector machines (SVMs). Univariate encoding models construct a relationship between experimental variables and the measured signal in one voxel which allows to statistically test activation differences between experimental conditions (Smith, 2004; Monti, 2011). Multivariate decoding algorithms extract experimental variables from the measured signals in many voxels which allows to reliably decode experimental conditions from brain activation (Haxby et al., 2001; Haxby, 2012; Cox and Savoy, 2003; Norman et al., 2006; Haynes and Rees, 2006; Haynes, 2015). This is commonly called “multivariate pattern analysis” (MVPA) for neuroimaging data.

MVPA for fMRI can either be performed by a single decoding from all measured signals in the entire brain (“whole-brain decoding”) or by several decodings from spatially well-circumscribed regions of interest (ROI; “ROI-based decoding”). Another, third option is given by building a sphere of voxels in some radius around each voxel and then decoding from measured signals in each of these “searchlights” separately (“searchlight-based decoding”) which gives a map of decoding performance for all searchlight positions across the brain. Searchlight-based decoding was introduced in the early days of MVPA to harness the statistical power of cross-validated prediction afforded by machine learning algorithms, but to also identify information at potentially unexpected locations in the brain (Kriegeskorte et al., 2006; Haynes et al., 2007). As such, searchlight decoding is not able to detect distributed activation patterns across multiple brain regions, but rather serves for localizing information in regional networks (Haynes, 2015). Moreover, the performance of searchlight decoding will depend on searchlight radius, size of the focal activation, information prevalence and other factors (Etzel et al., 2013).

Cross-validated prediction can either be performed over parameter estimates calculated from fMRI recording sessions (so-called “run-wise betas”) or it can be used to decode the identity of individual trials (decoding based on “trial-wise betas”). For trial-wise decoding, a common approach is to estimate trial-wise response amplitudes from the fMRI signal using a GLM with one onset regressor per trial (Rissman et al., 2004; Mennes et al., 2013), generated by convolution with a hemodynamic response function (HRF; Friston et al., 1998; Henson et al., 2001). This approach is called the “least squares, all” method (LS-A) and typically performs poorly for rapid event-related fMRI: When inter-trial-intervals are very short, the HRF regressors overlap in time due to the temporally extended shape of the canonical HRF, causing trial-wise estimates to be serially correlated and highly variable (Mumford et al., 2012; Turner et al., 2012) which can distort parameter estimates and invalidate statistical tests (Mumford et al., 2014).

One currently accepted approach for solving this problem is to estimate each trial’s response via a GLM including a regressor for that trial and another regressor for all other trials (Mumford et al., 2012). This approach is called the “least squares, separate” method (LS-S) and was found to outperform the LS-A method as well as a range of other techniques (Mumford et al., 2012, Fig. 3). The rationale behind LS-S is that the one-trial regressor is only weakly correlated to the all-other-trials regressor which effectively reduces the variance and auto-correlation of the trial-wise parameter estimates (Mumford et al., 2014). One disadvantage of LS-S is that each trial requires fitting a separate GLM, so that e.g. calculating activation patterns for 100 trials needs 100 GLMs.

**Figure 1:**
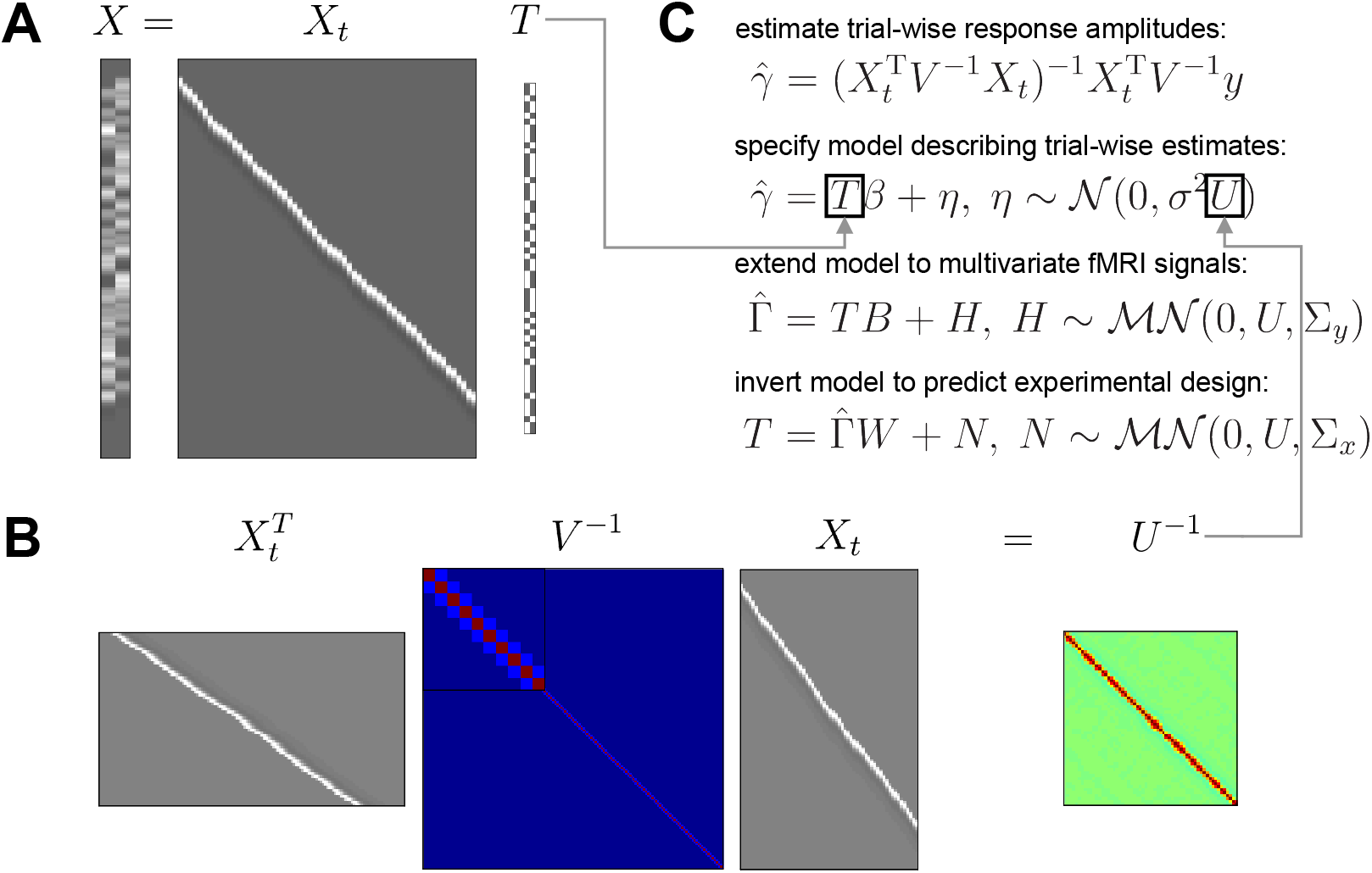
Mathematics of ITEM analysis. **(A)** The trial-wise design matrix *X*_*t*_ (scans *×* trials) can be related to the standard design matrix *X* (scans *×* conditions) using a trial-level specification matrix *T* (trials *×* condition). In this simple case, *T* is just an indicator matrix specifying which trial belongs to which condition. **(B)** Under this relationship, the (inverse of the) trial-by-trial covariance matrix *U* (trials *×* trials) is equal to the trial-wise design matrix *X*_*t*_, multiplied with itself and weighted by the scan-by-scan covariance matrix *V* (scans *×* scans). In the present case, overlapping HRFs induce serial correlation between temporally nearby trials. **(C)** Given that trial-wise response amplitudes *γ* are estimated from the voxel-wise fMRI signal *y* using the trial-wise design matrix *X*_*t*_ (first equation), it can be shown that they follow a linear model with the transformation matrix *T* as its design matrix and the uncorrelation matrix *U* as its temporal covariance (second equation). Combining the trial-wise parameter estimates from multiple voxels into a matrix 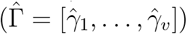 leads to a multivariate version of this model (third equation) and assuming a correspondence between forward and backward model (*BW* = *I*_*p*_) leads to an inverted version of this model (fourth equation).

**Figure 2:**
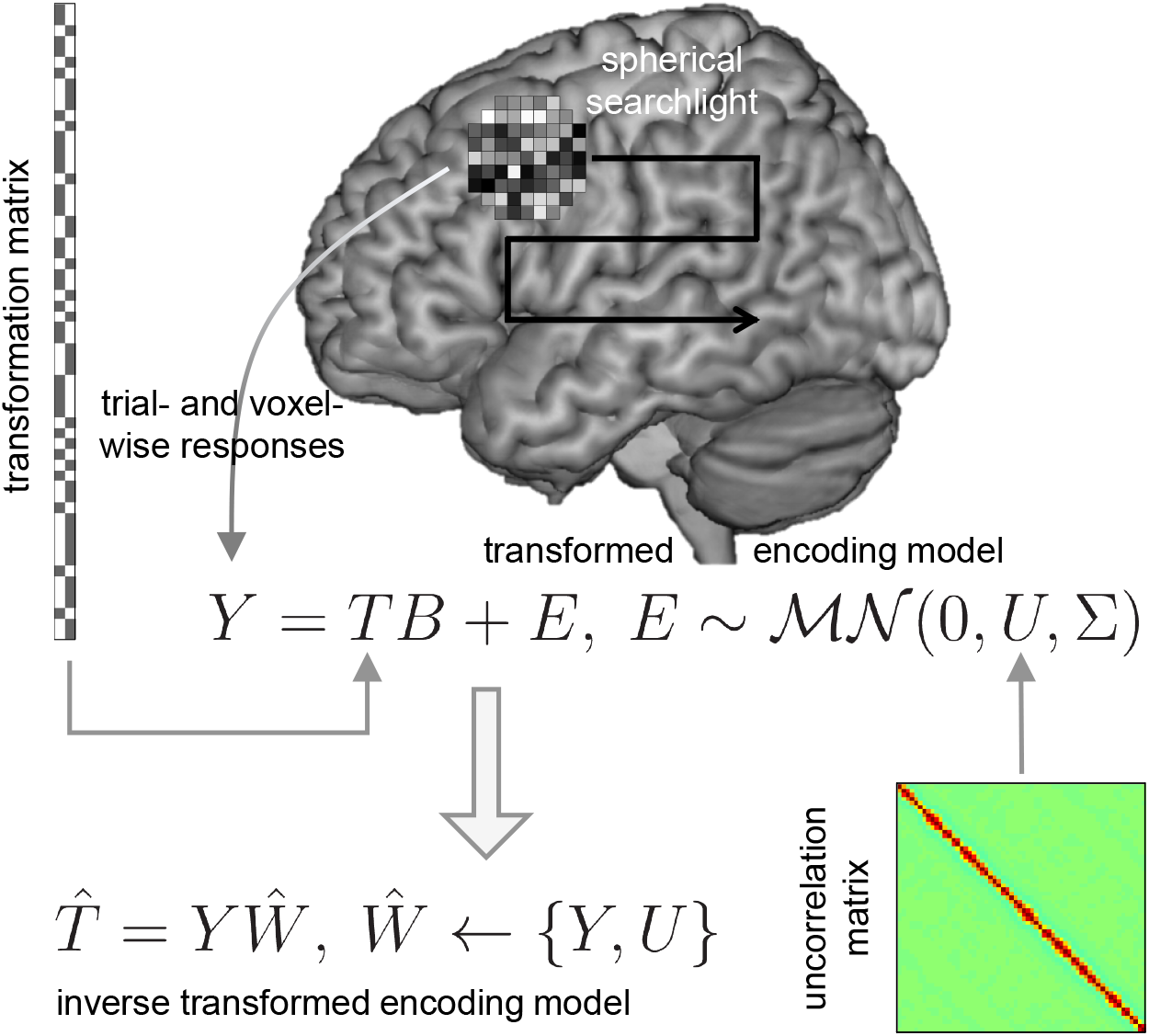
Searchlight-based ITEM analysis. The trial-wise parameter estimates *Y* (trials *×*voxels) are extracted from all voxels belonging to a spherical searchlight which is successively centered on each voxel in the brain (black arrow). The estimated responses *Y* follow a multivariate linear model which uses the transformation matrix *T* (see Figure 1A) as design matrix and the uncorrelation matrix *U* (see Figure 1B) as temporal covariance (gray arrows; for simplicity, 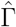 from Figure 1C is here replaced by *Y* and Σ_*y*_ from Figure 1C is here replaced by Σ). This is called a “transformed encoding model” (center equation). Such a model can be inverted by predicting experimental design variables *T* from the estimated responses *Y* and their covariance *U* via an estimated weight matrix *Ŵ* . This is called “inverse transformed encoding modelling” (bottom-left equation).

**Figure 3:**
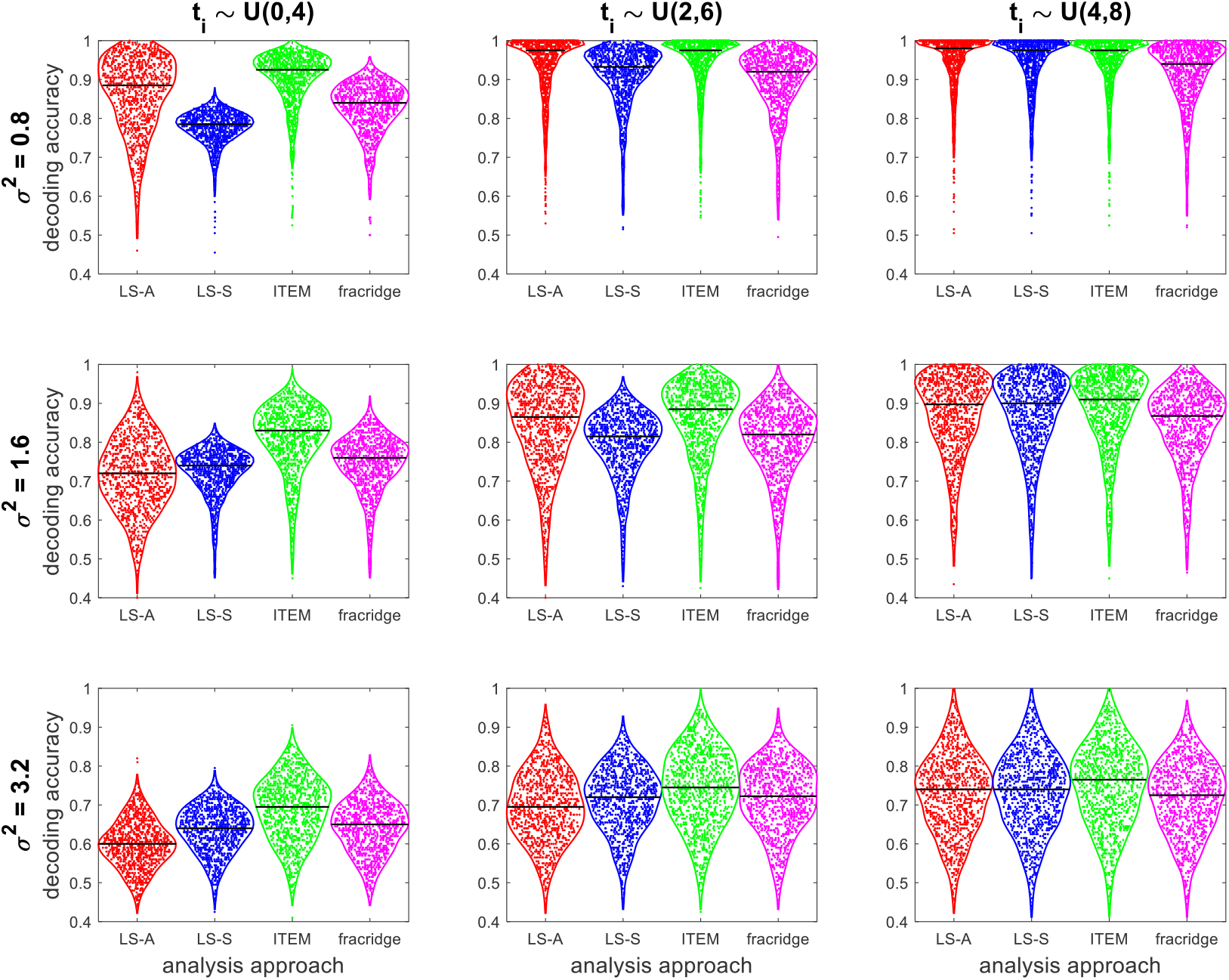
Simulation validation of multi-voxel ITEM analysis. For each combination of inter-stimulus-intervals (*t*_*i*_) and noise variance (*σ*^2^), decoding accuracies for classifying two experimental conditions (*N* = 1000 simulations) are given for LS-A (red), LS-S (blue), ITEM (green; the proposed approach) and GLMsingle based on fractional ridge regression (magenta). For long *t*_*i*_ and low *σ*^2^ (upper right), average decoding accuracies of all algorithms are close to 1. When the noise variance is high (bottom row) or inter-stimulus-intervals are short (left column), the ITEM approach outperforms LS-S and GLMsingle. In each violin plot, the horizontal black line denotes the median. For details of the simulation study, see Appendix A.

More recently, GLMsingle has been proposed as a new technique for improving the accuracy of single-trial response estimates (Prince et al., 2022). GLMsingle combines three advances in parameter estimation: First, each voxel receives a custom HRF from a library of candidate HRFs; second, noise regressors are estimated from fMRI time series using cross-validation across runs (Kay et al., 2013); and third, the variance of beta estimates is reduced via regularization using fractional ridge regression (Rokem and Kay, 2020). GLMsingle has been found to outperform LS-S for different data sets (Prince et al., 2022, Fig. 3), but it also increases the computational cost: GLMsingle estimation using the above-mentioned standard settings typically takes even longer than LS-S estimation.

In previous work, we have suggested “inverse transformed encoding models” (ITEM; Soch et al., 2020), a trial-wise modelling framework that builds on LS-A estimates, but accounts for the correlation between trials by incorporating their covariance matrix into a linear model operating at the trial level. Whereas LS-S estimates each trial’s response using a separate model (Mumford et al., 2012) and GLMsingle uses ridge-regularized, cross-validated single-trial parameter estimation, with optional HRF optimization and data-driven nuisance regressors (Prince et al., 2022), ITEM analytically incorporates the trial-by-trial covariance structure into decoding analysis. ITEM does not require fitting a separate GLM for each trial, thus extremely lowering the computational cost compared with LS-S. GLMsingle also fits a single GLM per run, but rather regularizes noisy dimensions while preserving the remaining variance.

Our first contribution was somewhat limited by the fact that (i) ITEM as a technique for multivariate decoding was validated using an univariate simulation (adapted from Mumford et al., 2012); and (ii) we used ROI-based decoding for a visual stimulation dataset (acquired by Heinzle et al., 2011) which did not explore the full potential of ITEM for localization of information in the brain. In this paper, we present and validate searchlight-based ITEM analysis (ITEM-SL), i.e. ITEM-style decoding from signals in spherical volumes of interest the centers of which are placed on each voxel in the brain. ITEM-SL here refers to the fact that an ITEM analysis is separately performed in each searchlight rather than just for a few regions of interest.

We perform a truly multivariate simulation in which we repeatedly generate data from synthetic searchlights and find that performance gains of ITEM-SL over LS-S are even higher than in our original simulation, suggesting that the advantage of the proposed approach over the popular LS-S approach grows with increasing number of voxels that are decoded from. Additionally, we apply ITEM-SL to a visual fMRI data set (Soch et al., 2023) and are able to recover known principles of visual cortex organisation, e.g. contralateral processing of the visual hemifields and polar-coordinate representation of the visual field. While LS-S and GLMsingle are also able to detect those patterns, ITEM-SL is computationally less expensive than the former two.

The structure of this paper is as follows. First, we will introduce ITEM-SL by recapitulating the theory behind ITEM-style analysis and describing the searchlight-based implementation (see Section 2 and Figure 2). Second, we will perform a simulation study on searchlight-based classification from synthetic fMRI data to demonstrate that ITEM is more powerful than combining the popular LS-S approach with SVM-based classification (see Section 3 and Figure 3). Third, we will describe an empirical application in which ITEM is used for searchlight-based reconstruction of local contrast at many visual field positions in an extremely rapid event-related design and thereby recover the well-known retinotopic mapping of early visual cortex (see Section 4 and Figure 5).

**Figure 4:**
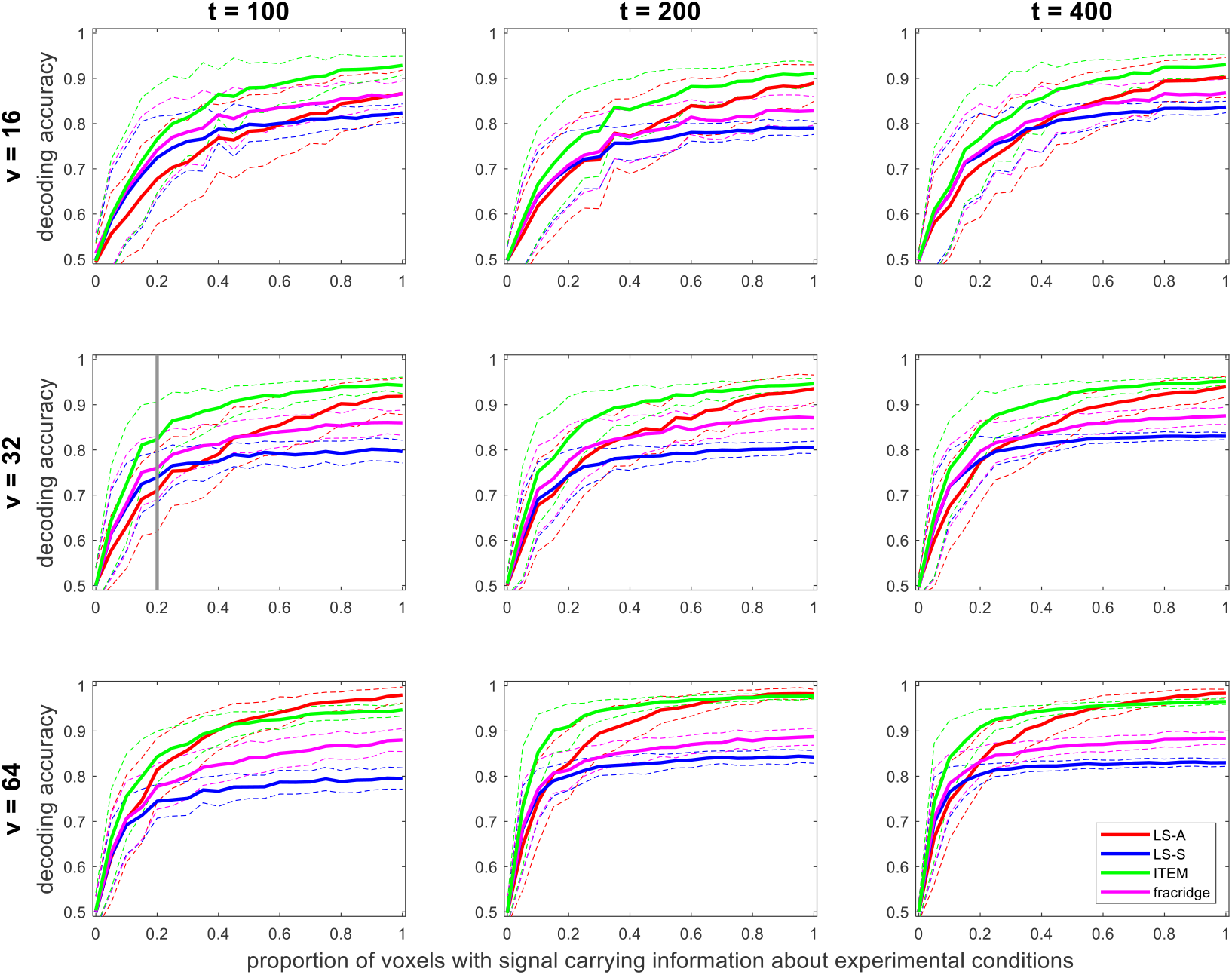
Effects of number of trials, voxels per searchlight and informative voxels. For each combination of number of trials (*t*) and number of voxels (*v*), decoding accuracies for classifying two experimental conditions are given as a function of the proportion of voxels (*r*) that carry information about the conditions (*N* = 100 simulations per value of *r*), separated between LS-A (red), LS-S (blue), ITEM (green; the proposed approach) and GLMsingle based on fractional ridge regression (magenta). Solid lines show average decoding accuracies, dashed lines correspond to one standard deviation. For *r* = 0, average decoding accuracies approach 0.5, and for *r* = 1, each method reaches its maximum average accuracy, given number of trials and voxels per searchlight. In the first panel in the second row, the scenario corresponding to our first simulation (see Figure 3) is highlighted with the vertical gray bar. For details of the simulation study, see Appendix A.

**Figure 5:**
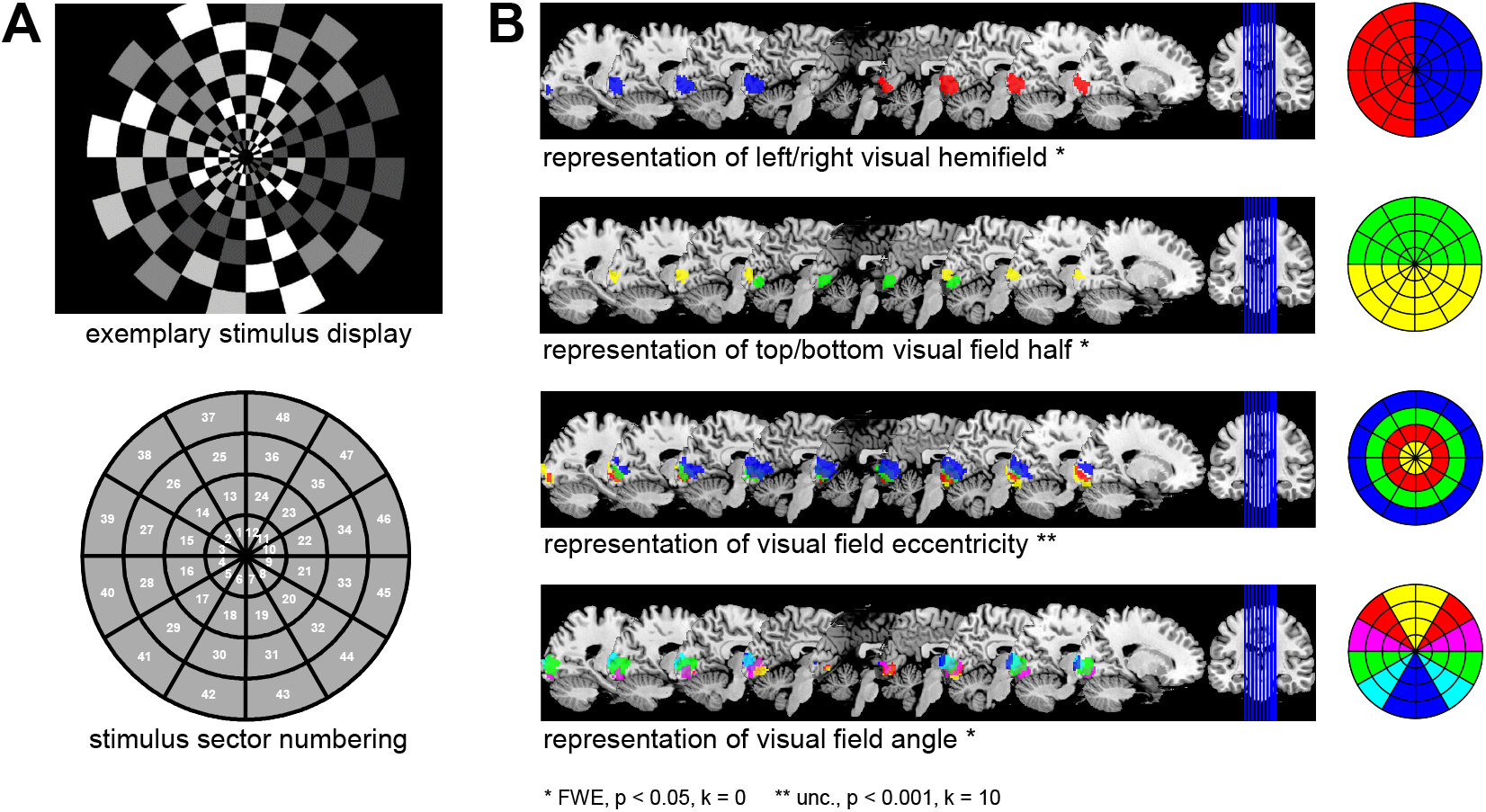
Empirical validation of searchlight-based ITEM analysis. **(A)** During fMRI scanning, subjects were stimulated with flickering checkerboard patterns (top) whose illumination intensity changed from trial to trial. The visual field was partitioned into 48 sectors (bottom) organized into 4 rings and 12 segments. Trial-wise intensity levels in all 48 sectors were reconstructed using ITEM-based searchlight decoding from searchlights centered on each voxel (SL radius = 6 mm) and predictive correlations between actual and reconstructed intensities were calculated for each searchlight. **(B)** Then, predictive correlation maps were normalized to standard space and submitted to a repeated-measures ANOVA with eccentricity (4 levels) and angular direction (12 levels) as within-subject factors. Colored voxels indicate searchlights from which the visual contrast in highlighted sectors could be decoded with average predictive correlation significantly greater than zero (unc. = uncorrected, FWE = family-wise error-corrected, *p* = significance level, *k* = voxel extent threshold).

## 2 Methods

In this section, we briefly summarize the methodology on which searchlight-based ITEM analysis is based. We review the problem of trial-by-trial correlations in fMRI decoding (see Section 2.1) and recapitulate how inverse transformed encoding models solve this problem (see Section 2.2). Then, we describe how an ITEM analysis works in practice (see Section 2.4) and explain the searchlight-based implementation of this approach (see Section 2.5). For more theory behind the methodology, see Soch et al., 2020.

### 2.1 Trial-wise decoding from fMRI

In univariate fMRI analysis, the goal is usually to investigate whether experimental conditions have statistically significant (or, significantly different) effects on measured responses in single voxels. These data are often analyzed using the “standard” general linear model (standard GLM)

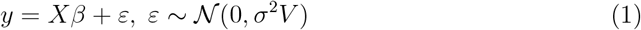

in which *y* is an *n ×* 1 vector of the measured BOLD signal in a single voxel (*n* = number of fMRI scans^1^), *X* is an *n × p* design matrix containing predictor variables (*p* = number of predictor variables, or “regressors”; Smith, 2004), *β* is a *p ×* 1 vector of regression coefficients and *ε* is an *n ×* 1 vector of error terms which are controlled by the noise variance *σ*^2^ and the *n × n* covariance matrix *V* .

In such analyses, *X* is typically known (because imposed by the experimental design) and *V* is estimated across all voxels (e.g. via restricted maximum likelihood; see Friston et al., 2002b; Friston et al., 2002a), but *β* and *σ*^2^ are unknown and have to be estimated for each voxel. Usually, *X* consists of “condition regressors”, i.e. trial onsets and durations convolved with the hemodynamic response function (HRF), and other regressors given as scan-by-scan covariates. Estimation of and inference based on (1) is usually referred to as the “mass-univariate approach” (Monti, 2011).

In multivariate fMRI analysis, the goal is sometimes to perform trial-wise decoding, i.e. to provide predictions for individual trials rather than collapsing them into conditions. In this case, it is advantageous to estimate trial-wise response amplitudes using a “trial-wise” general linear model (trial-wise GLM)

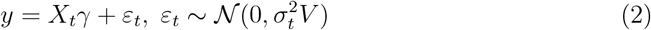

where *X*_*t*_ is an *n × t* design matrix with one HRF onset regressor for each trial (*t* = number of trials), instead of one such regressors for each condition, as in *X* (see Figure 1A); *γ* is a *t ×* 1 vector of trial-wise response amplitudes, sometimes also referred to as the “trial-wise betas” (cf. Rissman et al., 2004, p. 752); *ε*_*t*_ and 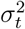 are the error terms and the noise variance of the trial-wise GLM, respectively.

Trial-wise response amplitudes can be estimated from (2) using e.g. weighted least squares which results in reponses commonly referred to as “LS-A estimates” (LS-A for “least squares, all”; Mumford et al., 2012)

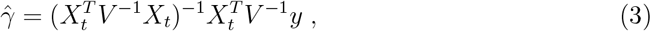

but this can become problematic: In rapid event-related designs, when inter-stimulus-intervals are short, the HRFs from adjacent trials can overlap in time, due to the comparably slow hemodynamic response with a peak at around 6 s and a post-stimulus undershoot until 20-30 s after stimulus onset (Friston et al., 1998). This induces serial correlations into the estimated trial-wise responses and makes those estimates more variable which reduces the statistical power of any analysis operating on them (Mumford et al., 2012; Turner et al., 2012).

For this reason, it has been proposed to estimate the response amplitude of each trial using a separate design matrix which leads to so-called “LS-S estimates” (LS-S for “least squares, separate”; Mumford et al., 2012)

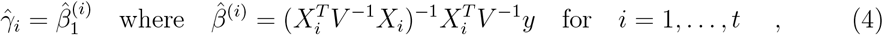

where *X*_*i*_ is an *n ×* 2 design matrix with one HRF regressor for the *i*-th trial and another regressor for all other trials. Thus, the *i*-th LS-S estimate is then given as the beta estimate corresponding to the first regressor of the *i*-th design matrix *X*_*i*_ (i.e. 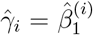) which requires that a separate GLM is run for each trial-wise parameter estimate. The rationale for this approach is that, because the second regressor contains *all other* trials, correlation with the first regressor is reduced which makes estimated trial-wise responses more robust. LS-S has been validated in previous work (Mumford et al., 2012; Turner et al., 2012; Mumford et al., 2014; Weeda, 2018) and is currently a widely used approach of extracting response estimates for trial-wise fMRI decoding.

### 2.2 Statistical theory behind ITEM analysis

Rather than artificially reducing the correlations between trial-wise parameter estimates which LS-S does, the ITEM approach attempts to naturally account for them by estimating and integrating their extent into the classification process. This starts by relating the design matrix *X* of the standard GLM to the design matrix *X*_*t*_ of the trial-wise GLM via a transformation matrix *T*

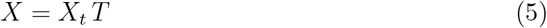

where *T* is a *t × p* matrix mapping from trials to conditions. This transformation matrix describes how individual trials combine into experimental conditions. In the simplest case, it is a binary matrix where *t*_*ij*_ = 1 indicates that trial *i* belongs to condition *j*, such that trial-wise HRFs are collected into onset regressors (see Figure 1A). However, *T* can also have continuous entries, such that trial-wise HRFs are linearly combined into parametric modulators and the corresponding column of *T* represents the values of the modulator variable (see Soch et al., 2020, Fig. 1B).

Upon making the assumption given by (5), it can be shown (see Soch et al., 2020, App. A) that the trial-wise parameter estimates from (3) follow a new linear model operating on the trial-by-trial level in which the design matrix is given by the transformation matrix *T* and the covariance matrix is a function of *X*_*t*_

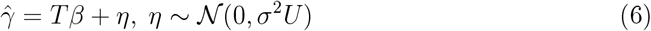

where 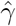 is a *t ×* 1 vector given by (3), *T* is the *t × p* matrix defined by (5) and *U* is a *t × t* matrix which specifies the trial-by-trial covariance and can be directly calculated from the trial-wise design-matrix^2^ (see Figure 1B):

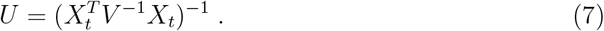

*U* is referred to as the “uncorrelation matrix”, because it allows to decorrelate trials and equation (6) is referred to as the “transformed encoding model”, because it operates on a transformed version of the measured data *y*, namely 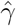. The transformed encoding model entails that estimated trial-wise responses linearly depend on experimental conditions and other design variables (embodied by *T* ) and that trial-by-trial correlations (embodied by *U* ) depend on the trial-wise design matrix used to estimate those responses.

Realizing that the response estimates from several voxels can be collected together, it extends into the “multivariate transformed encoding model” (MTEM; see Figure 1C)

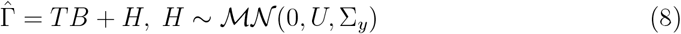

where 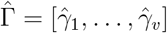 is a *t × v* matrix of trial-wise parameter estimates in a set of voxels (*v* = number of voxels) and Σ_*y*_ is a *v × v* matrix describing the spatial covariance, i.e. correlations in the activity of nearby voxels. By moving from (6) to (8), we simply combine many univariate into one multivariate linear regression model. Doing so, we are able to simultaneously model estimated trial-wise responses from many voxels by additionally accounting for the voxel-by-voxel correlations.

Finally, this model can be inverted to be turned into an “inverse transformed encoding model” (ITEM; see Soch et al., 2020, App. C)

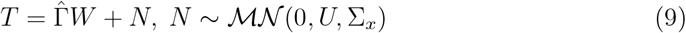

where *W* is a weight matrix mapping from trial-wise response amplitudes 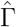 to the transformation matrix *T*, defined as the inverse of the activation pattern *B* via *B W* = *I*_*p*_; which implies that 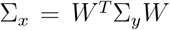 is a *p × p* matrix describing the ensuing condition-by-condition covariance of the ITEM^3,4^. Note that the matrix *T* contains experimental design variables, such as the experimental condition of each trial or associated modulator variables. This means that, using the inverse transformed encoding model, such design variables can be reconstructed (i.e. classified or regressed) from the estimated trial-wise responses 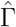 in the set of voxels considered.

### 2.3 Cross-validated prediction in ITEM analysis

In an ITEM analysis, the goal to estimate (9) in a cross-validated fashion in order to come up with a prediction for *T* . To this end, the data set is partitioned into fMRI recording sessions, and in each cycle, the training set is given by all but one session (e.g. runs 2-4) and the test set is given by the remaining session (e.g. run 1).

Given this partition, the weight matrix *W* is estimated from the trial-wise responses 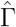 and the trial-by-trial correlations *U* belonging to the training data

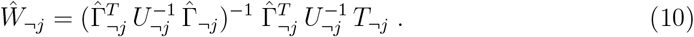

where *¬j* denotes all sessions except *j*. This estimate is then used to predict the trial-by-trial experimental design variables *T* for the test data

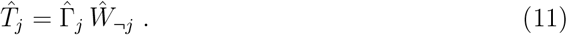

Based on the estimated transformation matrix 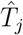 in the left-out session *j*, one can then perform classification between experimental conditions or make a regression of some target variable (see Section 2.5). This is repeated, until each session has once served as the test data (“leave-one-session-out cross-validation”).

### 2.4 Practical steps of an ITEM analysis

In practice, an ITEM analysis will proceed as follows:

1. *custom fMRI preprocessing*: Before any statistical analysis, fMRI data are preprocessed. Preprocessing is completely independent from ITEM-style analysis and can therefore be performed according to the preferences of the individual researcher.
2. *standard GLM analysis*: Then, a standard GLM analysis is performed. This is performed in the exact same way as for univariate fMRI analysis, i.e. using a condition-based, not trial-wise GLM, and can be done via standard packages, e.g. Statistical Parametric Mapping, Version 12 (SPM12; Ashburner et al., 2021).^5^
3. *trial-wise GLM estimation*: Then, based on this design information, a trial-wise design matrix is specified and trial-wise response amplitudes are estimated via (3). The ITEM toolbox for SPM (see Section 6.2) allows to distinguish conditions that are broken up into trials (e.g. target stimuli) vs. not broken up into trials (e.g. cue stimuli).
4. *trial-wise fMRI decoding*: Next, the actual predictive analysis is performed. In this step, trial-wise parameter estimates from a number of voxels (e.g. from a region of interest (ROI); or within searchlights (SL), see Section 2.5) are loaded and the desired decoding operation is performed using ITEM-style inversion of the model given by (9). The ITEM toolbox for SPM allows to select between
  a. classification (= decoding two or more experimental conditions) vs. regression (= decoding the values of a continuous target variable); as well as
  b. ROI-based decoding (= decoding from signals in a pre-specified region) vs. searchlight-based decoding (= decoding from signals within spheres on each voxel).
5. *group-level analysis*: If applicable, decoding results can be generalized to the population by integrating single performance values from ROI-based decoding or voxel-wise performance maps from SL-based decoding across subjects.

In the later empirical validation (see Section 4), we describe for each of these five steps, how our exemplary ITEM analysis was conducted.

### 2.5 Searchlight-based implementation

While trial-wise parameter estimation works on a voxel-wise level^6^, trial-wise fMRI decoding was previously only implemented as ROI-based analysis^7^. With this work, we provide searchlight-based implementations of ITEM-style analyses^8^ and also perform a comprehensive application of searchlight decoding (see Section 4).

Searchlight-based ITEM analysis consists of the following steps: First, the desired search-light radius is used to generate searchlights by taking each in-mask voxel as the center voxel, placing a sphere with the given radius around it and including all in-mask voxels inside the sphere (see Figure 2, top-right). Second, the trial- and voxel-wise responses 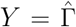 are extracted from each searchlight. For example, if the number of trials is 120 and the number of voxels per searchlight is 50, then *Y* will be a 120 × 50 signal matrix. Third, the transformation matrix *T* and uncorrelation matrix *U* are gathered to specify the transformed encoding model for each fMRI recording session (see Figure 2, center). Fourth, this model is inverted and variables of interest *T* are predicted via cross-validated estimation of the inverted model (see Figure 2, bottom-left).

Finally, if this has been repeated for all searchlights, a measure of decoding performance is calculated at each voxel by comparing the actual against the predicted values of the experimental design variables, across sessions:

- For *classification*, the column in the estimated matrix 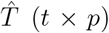 with the highest value is selected as the predicted condition in each trial, i.e. for each row. Then, decoding performance is calculated as decoding accuracy, i.e. the number of correct classifications, divided by the total number of trials.
- For *regression*, a column in the estimated matrix 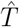 is taken as the set of predicted target values for this regressor. Then, decoding performance is calculated as the correlation between the predicted and actual values for this regressor.
- Note that ITEM outputs trial-wise predicted target values for all searchlights, such that the user is free to calculate other performance measures for discrete prediction, e.g. balanced accuracy (BA; Brodersen et al., 2010), or continuous prediction, e.g. median absolute error (MAE; Poldrack et al., 2020).

Decoding performances are stored in a single map for (a) each contrast to classify or (b) each regressor to predict and can later be used for voxel-wise group-level analysis.

### 2.6 Comparison with GLMsingle

GLMsingle is another technique for estimating trial-wise response amplitudes for later multivariate pattern analysis (Prince et al., 2022), based on a custom HRF for each voxel, cross-validated estimation of noise regressors (Kay et al., 2013) and fractional ridge regression (Rokem and Kay, 2020). GLMsingle is a natural competitor of ITEM, but there are a number of differences that we want to highlight:

- GLMsingle uses binary design matrices as input, such that trial onset and durations must be given in units of whole TRs.^9^ For designs that are not scan-locked (which is quite common in higher-cognitive function experiments), GLMsingle therefore approximates trial-wise parameter estimates. When event onsets do not exactly align with fMRI acquisition, one solution is to resample or upsample measured signals to the desired temporal resolution.^10^ Due to the slow nature of the hemodynamic response, these corrections are likely to capture meaningful temporal variations.
- Both, ITEM and GLMsingle accept user-supplied nuisance regressors (e.g. motion parameters or global signals). ITEM is able to assess the covariation between condition/trial-evoked responses and nuisance-related signals via its use of the uncorrelation matrix (see eq. 7; see Soch et al., 2020, Fig. 2B). Going beyond user-supplied regressors, GLM-single also offers the option to include data-driven nuisance regressors via GLMdenoise, which can capture further spatially correlated noise components.
- ITEM is designed to work with SPM.mat as input, restricting its use to SPM running in MATLAB. In contrast to that, GLMsingle is not connected to a particular software package, but can be used in both, MATLAB and Python (Prince et al., 2022), which offers more flexibility to users of platforms other than SPM.

In sum, we have included GLMsingle (alongside with LS-A, LS-S and ITEM) into our simulation studies (see Section 3) as well as the empirical analysis of a continuous visual stimulation experiment (see Section 4). We found that ITEM outperforms GLMsingle in simulations (see Figure 3) whereas performance differences for real data are negligible, ITEM and GLMsingle performing about equally well (see Figure 6).

**Figure 6:**
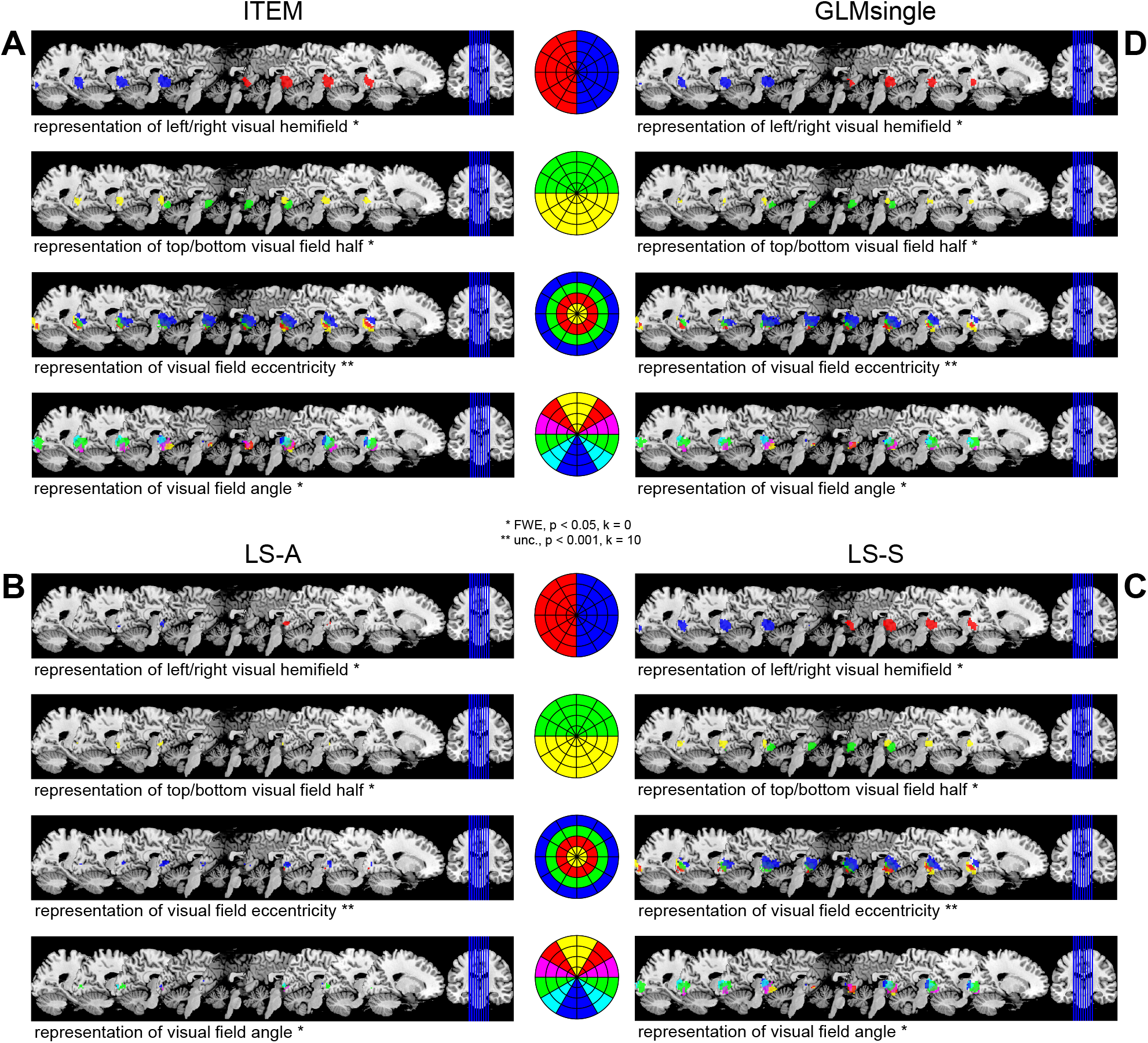
Comparison of searchlight-based ITEM with alternative approaches. The results of **(A)** searchlight-based ITEM analysis (identical with Figure 5B) are compared with results obatined when applying searchlight-based support vector regression to **(B)** voxel-wise LS-A estimates, **(C)** voxel-wise LS-S estimates and **(D)** voxel-wise estimates obtained with GLMsingle. The group-level analysis is identical for all approaches (see Figure 5). Colored voxels indicate searchlights from which the visual contrast in highlighted sectors could be decoded with average predictive correlation significantly greater than zero (unc. = uncorrected, FWE = family-wise error-corrected, *p* = significance level, *k* = voxel extent threshold).

## 3 Simulation

To validate searchlight-based ITEM using synthetic data, we adapt a simulation from Soch et al., 2020 which was itself adapted from Mumford et al., 2012. The main change was replacing the previously univariate generative model by multivariate signals generated in the present simulation. All simulation code is available from GitHub (see Section 6.2).

### 3.1 Methods

In our simulation, we compare four approaches of trial-wise decoding from fMRI signals introduced earlier: the naïve approach operating on uncorrected trial-wise parameter estimates (Mumford: “least squares, all”, LS-A); a popular approach based on parameter estimation using separate models (Mumford: “least squares, separate”, LS-S); the new approach proposed here, i.e. searchlight-based inverse transformed encoding modelling (ITEM-SL); and an alternative approach, GLMsingle.

LS-A entails decoding without accounting for correlation and taking trial-wise parameter estimates 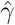 from equation (3) “as is”. LS-S is based on trial-wise parameter estimates based on equation (4), using a separate design matrix *X*_*i*_ for each trial *i* = 1, …, *t*, including one regressor for this trial and one regressor for all other trials. ITEM uses the same estimates as LS-A, but accounts for their correlation by incorporating the trial-by-trial covariance matrix *U* as in equation (9). GLMsingle performs regularized ridge regression to estimate trial-wise responses (see Section 2.6).

In the simulation, data were generated as follows: First, trials were randomly sampled from two experimental conditions, A and B. Second, voxel-wise average responses *µ*_*A*,*j*_ and *µ*_*B*,*j*_ (*j* indexes voxel) were sampled from the standard normal distribution *N* (0, 1). Third, trial-wise response amplitudes *γ*_*i*,*j*_ (*i* indexes trials) were sampled from normal distributions 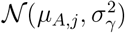 and 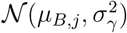 where *σ*_*γ*_ = 0.5.

Fourth, inter-stimulus-intervals *t*_*i*_ were sampled from uniform distributions 𝒰(0, 4) or𝒰(2, 6) or 𝒰(4, 8). Fifth, the design matrix *X*_*t*_ was generated based on the *t*_*i*_’s and convolution with the canonical HRF using stimulus duration *t*_dur_ = 2 s and repetition time TR = 2 s. An example design matrix for the case *t*_*i*_ *∼𝒰* (0, 4) is given in the middle of Figure 1A. Finally, a multivariate signal was generated by multiplying the trial-wise design matrix *X*_*t*_ with trial-wise response amplitudes Γ and adding zero-mean Gaussian noise *E* with variance *σ*^2^ where *σ*^2^ *∈ {*0.8, 1.6, 3.2*}* .

This was repeated for *N* = 1,000 simulations with *S* = 2 sessions and *t* = 100 trials per session (50 per condition). Each simulation can be seen as an individual searchlight and the number of voxels per simulation/searchlight was *v* = 33 which corresponds to a spherical searchlight with a radius of 2 voxels^11^. A proportion of voxels with information *r* was specified as *r* = 20%, such that for 1 − *r* = 80% of the voxels, *µ*_*B*,*j*_ was set to *µ*_*A*,*j*_, implying no difference between A and B in those voxels. A more detailed description of the simulation study is given in Appendix A.

For LS-A and ITEM, trial-wise parameter estimates 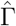 were obtained by least-squares estimation using equation (3). For ITEM, 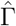 was subjected to an additional restricted maximum likelihood (ReML) analysis (see Soch et al., 2020, App. B), in order to separate the natural trial-to-trial variability (coming from 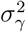) from the induced trial-by-trial correlations (coming from *X*_*t*_). For LS-S, 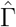 was obtained via trial-specific design matrices *X*_*i*_ (*i* = 1, …, *t*) using equation (4). For GLMsingle, trial-wise parameter estimates were obtained using a simplified configuration of the GLMsingle toolbox, with HRF optimization and GLMdenoise disabled, focusing solely on fractional ridge regression (fracridge). Note that this is different from the full GLMsingle pipeline which includes HRF selection and data-driven noise regressors (Prince et al., 2022).

In a second simulation, we focused on one of these simulation scenarios – characterized by short inter-stimulus-intervals *t*_*i*_ *∼ 𝒰* (0, 4) and medium noise variance *σ*^2^ = 1.6 – and investigated the effects of the number of voxels per searchlight *v ∈ {*16, 32, 64*}*, the number of trials per session *t ∈ {*100, 200, 400*}* and the proportion of voxels with information 0 *≤r ≤*1. For each combination of number of voxels *v*, number of trials *t* and proportion of voxels *r*, decoding accuracies of LS-A, LS-S, ITEM and GLMsingle were compared across *N* = 100 simulations.

### 3.2 Results

Given that there are differences between them, the two experimental conditions can be decoded from the generated data using a classification algorithm. For this purpose, we here chose support vector machines (SVM) for classification (SVC). For LS-A, LS-S and GLMsingle, condition labels for A and B are coded as 1 and 2 and the corresponding SVM is calibrated based on one session. Then, condition labels are predicted from trial-wise response amplitudes in the left-out session. For GLMsingle, trial-wise parameter estimates were scaled inside each voxel (subtracting the mean, dividing by standard deviation) to account for magnitude differences between estimates across voxels.

For ITEM, as trial-by-trial correlations cannot easily be accounted for by SVC, the linear decoding procedure outlined in Section 2 was employed for cross-validated classification of trial types. Trial-wise decoding was performed by predicting experimental design information using equations (10) and (11). For all approaches, decoding accuracy (DA), i.e. the percentage of trials correctly assigned across both sessions, was used as the measure of decoding performance. Each procedure leads to one DA value per simulation/searchlight, the distributions of which are visualized as violin plots.

In our first simulation, there is a real effect in *r* = 20% of the voxels. In each simulation scenario, ITEM outperforms LS-S by between 0.0% (*σ*^2^ = 0.8, *t*_*i*_ *∼ 𝒰* (4, 8)) and 14.0% (*σ*^2^ = 0.8, *t*_*i*_ *∼ 𝒰* (0, 4)) in terms of median decoding accuracy (see Figure 3). Also, ITEM outperforms GLMsingle by between 2.3% (*σ*^2^ = 3.2, *t*_*i*_ *∼ 𝒰* (2, 6)) and 8.5% (*σ*^2^ = 0.8, *t*_*i*_ *∼ 𝒰* (0, 4)). This is particularly the case when inter-stimulus-intervals are short (all *σ*^2^ for *t*_*i*_ *∼ 𝒰* (0, 4)). Note that even LS-A outperforms LS-S and GLMsingle for low-variance situations (all *t*_*i*_ for *σ*^2^ = 0.8). In conclusion, ITEM outperforms alternative approaches in terms of sensitivity, for the simulation scenarios considered here.

In our second simulation, we investigated the effects of number of trials *t*, number of voxels *v* and proportion of voxels with information about experimental conditions *r*, on decoding accuracy, for one scenario from the first simulation (*σ*^2^ = 1.6, *t*_*i*_ *∼ 𝒰* (0, 4)). As expected, we find that decoding performance mildly increases with more trials (cf. third vs. first column of Figure 4) and more strongly increases with more voxels (cf. third vs. first row of Figure 4). Moreover, when setting the proportion of activated voxels to *r* = 0%, such that no difference between the conditions exists, all approaches considered have an average decoding accuracy of around 50% (see Figure 4). With *r >* 0%, such that conditions differ, ITEM’s average decoding accuracy is generally above that of LS-A, LS-S and GLMsingle. With increasing proportion of selective voxels, average decoding accuracy goes up, but surprisingly, LS-S and GLMsingle show much weaker performance than LS-A for voxel proportions close to *r* = 100% (see Figure 4).

## 4 Application

To validate searchlight-based ITEM using empirical data, we analyze fMRI data that were acquired by Heinzle et al., 2011 and are more closely described in Soch et al., 2023. This data set was originally acquired to investigate relationships between sensory-visual and cortico-cortical receptive fields and is here used to recover the spatial organization of early visual cortex. The entire data set is available from OpenNeuro (see Section 6.2).

### 4.1 Experiment

Because the descriptor of this data set is available open access (Soch et al., 2023)^12^, the experimental design is reported rather shortly in this section.

Four right-handed, healthy subjects (24-28 years, 3 male, 1 female) participated in a visual stimulation experiment in which they viewed a dartboard-shaped stimulus that consisted of 48 “sectors” organized into 4 “rings” and 12 “segments” (see Figure 5A). Each sector was a flickering checkerboard randomly changing its local visual contrast every 3 s across 8 recording sessions with 100 trials per session.

Intensity levels were logarithmically spaced between 0.1 and 1 and used for analysis as linearly spaced between 0 and 1 in steps of 1/3. Importantly, there was no inter-stimulus-interval, implying the maximum possible overlap between HRFs at this stimulus duration and constituting a perfect application case of the presented approach of trial-wise decoding in the presence of trial-by-trial correlations.

To maintain fixation at the center of the visual display, subjects were engaged in a cognitive control task. Landolt’s C was presented in the middle of the screen, and subjects had to indicate whether it opened to left or to the right side.

Functional magnetic resonance imaging (MRI) data were collected on a 3-T Siemens Trio with a 12-channel head coil. In each session of the visual stimulation experiment, 220 T2*-weighted, gradient-echo EPIs were acquired at a repetition time TR = 1500 ms, echo time TE = 30 ms, flip-*α* = 90° in 25 slices (slice thickness: 2 mm (+1 mm gap); matrix size: 64 *×* 64) resulting in a voxel size of 3 *×* 3 *×* 3 `mm.`

### 4.2 Analysis

The five steps of ITEM analysis (see Section 2.4) for these data were as follows:

1. *custom fMRI preprocessing*: Data were converted to the BIDS format (Gorgolewski et al., 2016), reoriented to the axis from commissura anterior (AC) to commissura posterior (PC), corrected for acquisition time (slice timing) and head motion (spatial realignment) using SPM12.
2. *standard GLM analysis*: The first-level design matrix included 1 “condition” regressor for continuous visual stimulation; 48 parametric modulators describing the intensity levels in all sectors; 2 regressors of no interest for the control fixation task; further nuisance regressors for movement parameters and temporal filter; and a constant regressor modelling the implicit baseline.
3. *trial-wise GLM estimation*: The ITEM toolbox function `ITEM_est_1st_lvl` was used to estimate trial-wise response amplitudes where only the continuous visual stimulation events (100 trials), but not the control fixation task events (Landolt’s C) was broken up into trial-wise structure, as our major interest was predicting local visual contrast from concurrent early visual cortex activity.
4. *trial-wise fMRI decoding*: The ITEM toolbox function `ITEM_dec_recon_SL` was used to perform searchlight-based regression of intensity levels in all sectors from searchlights all over the brain using a searchlight radius of 6 `mm` to yield a correlation coefficient (CC) map for each sector and subject.
5. *group-level analysis*: Finally, CC maps were normalized into the common MNI space and subjected to a repeated-measures ANOVA with visual field radius (4 rings = 4 levels) and visual field angle (12 segments = 12 levels) as within-subject factors. Using suitable contrasts, we were looking for voxels in which the average CC for a subset of sectors was significantly larger than zero (see Figure 5B).

In addition to searchlight-based ITEM analysis and in order to compare it against the existing alternatives, fMRI decoding was also performed using the LS-A approach, the LS-S approach (Mumford et al., 2012) and with GLMsingle (Prince et al., 2022). For these analyses, steps 1, 2 and 5 were identical and steps 3 and 4 were modified as follows:

- *trial-wise estimation*: Custom MATLAB code was used to obtain voxel-wise LS-A estimates according to equation (3) and voxel-wise LS-S estimates according to equation (4). The GLMsingle toolbox was used to obtain voxel-wise paramater estimates with an HRF tailored to each voxel, with GLMdenoise regressors, with movement parameters from spatial realignment as extra nuisance regressors and with ridge regression regularization (“type-D model” outputs).
- *trial-wise decoding*: LS-A, LS-S and GLMsingle parameter estimates were subjected to support vector regression (SVR) for predicting each sector’s intensity levels for all trials, employing searchlights of radius 6 `mm` and leave-one-session-out cross-validation. Actual and predicted values were compared with Pearson correlation, resulting in one correlation coefficient (CC) map for each sector and subject.

The complete empirical data analysis can be reproduced using MATLAB code available from GitHub (https://github.com/JoramSoch/ITEM-SL-paper).

### 4.3 Results

Results obtained from the repeated-measures ANOVA matched well-known properties of early visual cortex (see Figure 5B): (i) contralateral processing of visual hemifield, i.e. left visual field activates right visual cortex and vice versa; (ii) representation of visual field half, i.e. top visual stimulation activates medial parts, bottom visual stimulation activates lateral parts of visual cortex; (iii) representation of eccentricity along a posterior-anterior axis, i.e. more outer parts activate more anterior regions; (iv) representation of angular direction along a dorsal-ventral axis, i.e. more bottom parts activate more dorsal regions; and (v) taking (iii) and (iv) together, polar-coordinate representation of the visual field in primary visual cortex (Zeidman et al., 2018).

All the results were significant at *α* = 0.05, whole-brain corrected for family-wise error; except for result (iii) for which uncorrected inference with a significance threshold *α* = 0.001 and an extent threshold *k* = 10 was applied (see Figure 5B).

When comparing searchlight-based ITEM with alternative approaches, we find a clear pattern for all these properties of early visual cortex: Whereas LS-A mostly fails to identify selectivity to visual field hemisphere, eccentricity and angle (see Figure 6B), results are qualitatively identical for ITEM, LS-S and GLMsingle (see Figure 6A/C/D), although quantitatively, there are less significant voxels when using GLMsingle (cf. Figure 6A vs. Figure 6D). This suggests that ITEM, LS-S and GLMsingle are about equally capable of identifying regularities in the human visual system from this data set.

## 5 Discussion

We have extended inverse transformed encoding models (ITEM), a previously proposed method for dealing with trial-by-trial correlations in fMRI decoding, from region-of-interest (ROI) to searchlight-based (SL) analysis. Whereas earlier contributions to trial-level prediction from fMRI responses have suggested ad-hoc solutions, e.g. estimating each trial’s response amplitude using a different model in order to reduce trial-by-trial correlations (Mumford et al., 2012, 2014), the present technique offers a principled approach by accounting for the actual distribution of trial-wise parameter estimates. By going beyond individual ROIs to decoding from the vicinity of each voxel in the brain (see Figure 2), we have demonstrated that SL-based ITEMs can be successfully used for information-based mapping (Kriegeskorte et al., 2006; Haynes et al., 2007) of cognitive functions, e.g. visual perception (see Figure 5).

The proposed approach is conceptually similar, but mathematically different from inverted encoding models (IEM; Brouwer and Heeger, 2009; Saproo and Serences, 2014; Gardner and Liu, 2019; Sprague et al., 2019). While IEMs are based on inverting an estimated forward model, ITEM consists in estimating a probabilistically derived inverse model. Another difference is that the IEM approach ignores covariance between trials which ITEMs are designed to capture. For an in-depth discussion, see our previous contribution on ITEM (see Soch et al., 2020, App. D).

### 5.1 Assessment of simulation validation

In our simulation study, we used two experimental conditions, because cognitive neuroscience experiments often comprise binary experimental designs and two-class comparisons are even used when there are more experimental conditions, e.g. to infer on the effect of a (two-level) factor in a factorial design, collapsing across the levels or separating between the levels of the respective other factor(s). On the other hand, there is no reason to believe that our simulation results do not hold for more experimental conditions or factors with more than two levels, subject to the constraint that the chance level will change (e.g. 0.25 for four conditions).

In our earlier study, results on the relative advantage of ITEM over LS-S, one currently popular approach, were somewhat inconclusive: Whereas simulation studies found that gains in statistical power and classification accuracy were rather marginal (mostly between 0 and +2%, minimally − 0.83%, maximally +8.33% in favor of ITEM; see Soch et al., 2020, Fig. 5), the empirical application resulted in significantly higher predictive correlations for the ITEM approach (*p <* 0.001; ITEM: mean *r* = 0.31; LS-S: mean *r* = 0.25; see Soch et al., 2020, Fig. 7).

**Figure 7:**
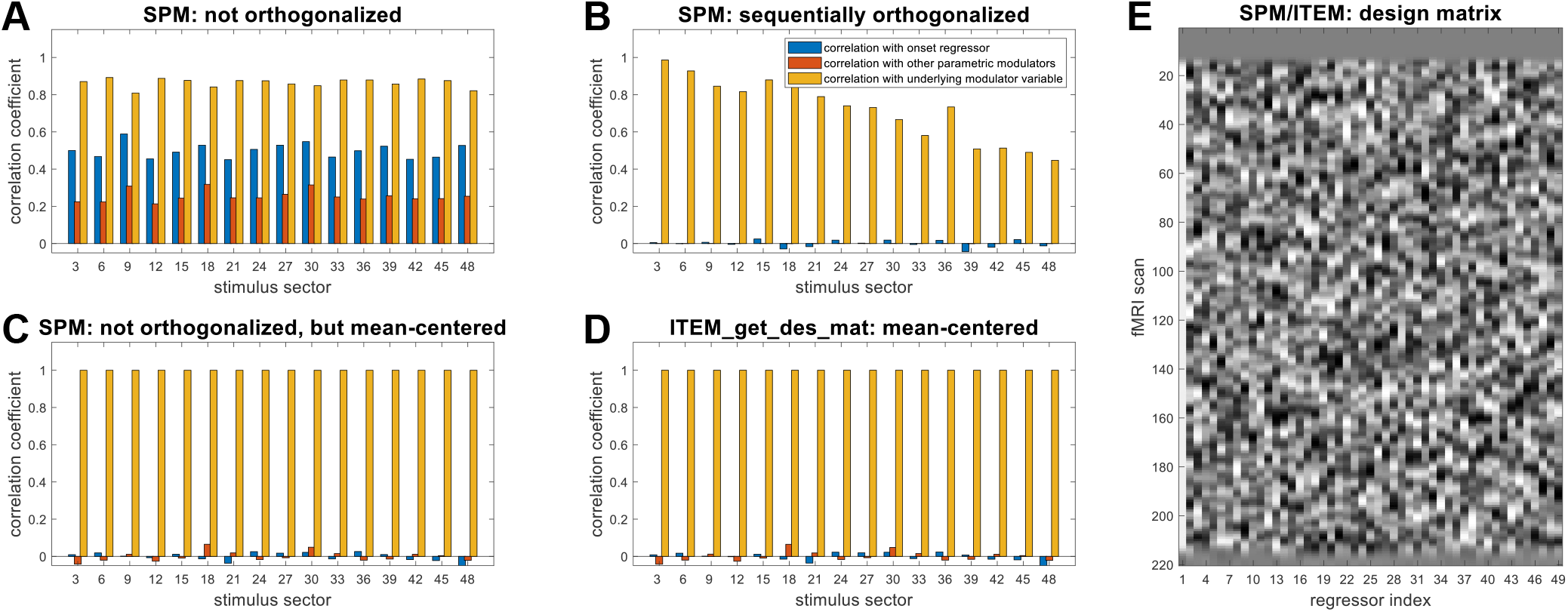
Effects of orthogonalization on parametric modulators. Pearson correlation co-efficients of all 48 parametric modulator regressors (i) with the onset regressor (blue), (ii) with the other parametric regressors (orange) and (iii) with the original modulator variables (yellow) **(A)** when using default SPM with orthogonalization switched off, **(B)** when using SPM’s sequential orthogonalization, **(C)** when using SPM without orthogonalization, but mean-centering beforehand and **(D)** when using the ITEM toolbox script for design matrix creation (which applies mean-centering, but not orthogonalization). For better readability of the figure, only every third parametric modulator is plotted. Note that not orthogonalizing or mean-centering induces correlations with the onset regressor (A, blue) and between the regressors (A, orange), but sequentially orthogonalizing makes parametric regressors become less and less faithful to the underlying modulator variables (B, yellow). **(E)** The design matrix that results from the procedure in C or D, without nuisance variables. Note that there are 49 regressors, the onset regressor for continuous stimulation (labeled as first regressor) and the parametric modulators for sector intensities (labeled with regressor index, if plotted in A-D).

We believe that this was due to the simulation study using only univariate signals, therefore producing only mild differences between methodologies (cf. LS-A in Soch et al., 2020, Fig. 5). Here, we have closed this gap and developed a truly multivariate (“searchlight”) simulation in which the ITEM approach outperformed or levelled with the LS-S method (always *≥* 0%, maximally +14% in favor of ITEM) and also outperformed the GLMsingle technique (always 2.3%, maximally +8.5% in favor of ITEM) in each simulation scenario considered (see Figure 3 and Section 3.2).

We hypothesize that the improvement of ITEM over LS-S with increasing number of voxels is due to the fact that the removal of temporal correlation is beneficial for each individual voxel. Thus, the more voxels the multivariate signal consists of, the higher the overall benefit in terms of decoding accuracy will be – also suggested by our second simulation actually varying the number of voxels (see Figure 4).

Also note that, like in the previous simulation study (see Soch et al., 2020, Fig. 5), we observed that with low error variance, LS-A outperforms LS-S (see Figure 3, top-left) – a pattern that continues to manifest when increasing the number of voxels and/or trials (see Figure 4, bottom-right). This suggests that the contamination of LS-A estimates with trial-by-trial correlations is not too harmful when the overall noise level is low, and LS-S (but not ITEM) might actually reduce statistical power compared to the naïve approach.

### 5.2 Assessment of empirical validation

In our earlier study, we applied ITEM in an ROI-based manner which required a feature selection step in which only voxels responsive to the task as such – no matter what aspect they are responsive to – were filtered out using a Bayesian model selection strategy (see Soch et al., 2020, p. 10). As a consequence of this, contrast levels in some sectors of the visual field could not be reconstructed with satisfactory precision – simply because the most task-responsive voxels exclusively represented visual receptive fields near the horizontal midline of the visual field (see Soch et al., 2020, Fig. 7D).

We believe that this was an unnecessary simplification and here replaced this ROI-based procedure by a searchlight-based approach (ITEM-SL) in which ITEM-style reconstruction was performed for multi-voxel trial-wise response amplitudes extracted from a spherical searchlight with radius *r* = 6 `mm` around each voxel inside the analysis mask. Following this, the fully parametric model accounting for covariation between all levels of angular direction and field radius was specified and estimated.

ITEM-SL showed very good sensitivity, as it reliably recovered the polar-coordinate organization of receptive field representation in primary visual cortex (see Figure 5B): When correcting for multiple comparisons at the whole-brain level, there were significant differences in reconstruction performance for visual hemifield, field half and field angle. For uncovering the representation of visual field eccentricity, uncorrected inference had to be applied (see Figure 5B, 1st/2nd/4th vs. 3rd row).

When using LS-A, LS-S (Mumford et al., 2012) or GLMsingle (Prince et al., 2022) as alternative approaches for trial-wise parameter estimation and combining them with searchlight-based SVR to decode local visual contrast in the same experiment, similar patterns for recovering the organisation of early visual cortex could be observed, except for LS-A (see Figure 6B/C/D), suggesting that LS-S, ITEM and GLMsingle can be regarded as state-of-the-art solutions delivering acceptable results.

ITEM-SL also exhibited high specificity, as no searchlights outside the occipital lobe could be used to decode visual contrast in low-level visual receptive fields and thus, no differences in reconstruction performance between visual field sectors were observed outside the visual cortex. Taken together, ITEM-SL can therefore be a powerful tool in the multivariate localization of cognitive functions in the human brain.

### 5.3 Assumptions and limitations

When applying a statistical technique to empirical data, it is important to keep in mind the assumptions made by this method. For ITEM-SL, the two most important assumptions are (i) linearity of the effects and (ii) normality of the errors^13^:

- The multivariate general linear model (MGLM) for fMRI assumes that the effects of the predictor variables (i.e. experimental conditions and modulator variables) on the measured variables (i.e. the measured BOLD signals) are linear in every voxel. For discrete experimental conditions, this means that average responses between conditions can differ. For continuous modulator variables, this means that average responses parametrically follow the levels of the parametric modulator. If there is no effect, this corresponds to a linear effect with a weight of zero.
- The MGLM further assumes that errors, i.e. additive parts of the measurements that cannot be explained by the predictor variables, are matrix-normally distributed with a fixed temporal covariance between trials that is derived from the trial-wise design matrix and an unknown spatial covariance between voxels which is unconstrained and fully estimated (see *𝒰* and Σ on Figure 2 or in Equation 8). Inside the searchlight, the voxel-by-voxel covariance is assumed to be constant over trials and the trial-by-trial covariance is assumed to be constant over voxels.

If any of the above assumptions is not met, the proposed technique should not be applied – or be applied with caution. Further research is necessary to assess how robust ITEM-SL is relative to violations of these assumptions.

For fMRI, there is good evidence that functional responses are linear relative to stimulation, especially in V1 (Boynton et al., 1996), and when the non-linear character of the hemodynamic response is accounted for (Friston et al., 1998). Furthermore, errors are usually assumed to follow normal distributions based on the central limit theorem and the rationale that every voxel’s signal represents a sum of a large number of physiological sources (Allefeld and Haynes, 2014).

Since the spatial covariance within a searchlight is largely determined by the physiological properties of nearby voxels, it is reasonable to assume that the voxel-by-voxel covariance is constant over trials. Moreover, since trial-wise response amplitudes are estimated using the same trial-wise HRF regressors for all voxels in a searchlight, it is reasonable to assume that the trial-by-trial covariance is constant over voxels. The most critical dependency of ITEM-SL is therefore whether the assumed trial-to-trial covariance structure holds. Our empirical validation provides grounds to believe this.

Something which ITEM-SL is not capable of is to capture (i) non-linearity in multivariate patterns, e.g. non-linear boundaries between experimental conditions or saturating effects of modulator variables, and (ii) non-stationarity in multivariate patterns, i.e. changes of neural responses or noise structure over time. In such a case, a statistical model explicitly accounting for such cases should be employed. For example, support vector machines allow for curved class boundaries using the kernel trick (Boser et al., 1992), but they are, without further extension, also limited to constant responses over time. Moreover, we want to emphasize that any other method, while possibly allowing for non-linearity or non-stationarity, will likely be limited in its ability to capture between-trial correlations – which is the critical part of the present contribution.

## 6 Statements

### 6.1 Ethics Statement

When acquiring the data set used in this study, written informed consent was obtained from all subjects before participating in the experiments (Heinzle et al., 2011; Soch et al., 2023). The study was approved by the ethics committee of the University of Leipzig, Germany and conducted according to the Declaration of Helsinki.

### 6.2 Data and Code

SPM12-compatible MATLAB code for searchlight-based ITEM classification and regression has been added to the ITEM toolbox^14^. All code underlying the analyses in this paper is also available from GitHub^15^.

The data set used for empirical validation in Section 4 has been BIDS-formatted and uploaded to OpenNeuro^16^. Further instructions on data processing can be found in the readme file of the accompanying repository^17^.

### 6.3 Author Contributions

JS: Conceptualization, Data curation, Formal analysis, Investigation, Methodology, Software, Validation, Visualization, Writing – original draft, Writing – review & editing.

## 6.6 Acknowledgements

The authors would like to thank Carsten Allefeld and John-Dylan Haynes for helpful discussions of the manuscript’s contents as well as Jakob Heinzle for acquiring the fMRI data set used for empirical validation.

## 7 Appendix

### A Detailed information for simulation study

The generative model underlying the first simulation can be described as follows. First, in each session from each simulation run, *t* = 100 trials are evenly distributed into 2 experimental conditions or trial types (tt)

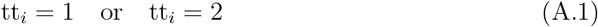

where tt_*i*_ = 1 and tt_*i*_ = 2 for 50 trials each.

Second, voxel-wise averages (i.e. average activity per condition and voxel, not trial-wise responses) are independently sampled from a standard normal distribution

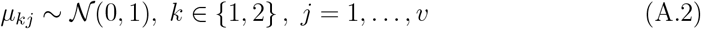

where *k* indexes trial type and *j* indexes voxel.

For a percentage of 1 − *r* = 80% voxels, activities were equalized between trial types

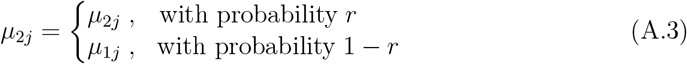

where the first line indicates that the activities sampled in (A.2) were kept. As the conditions are thus different with probability *r*, this value may be seen as the proportion of voxels with information about the conditions.

Third, trial-wise responses are independently sampled from a normal distribution

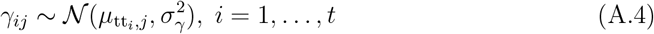

where *i* indexes trial and *σ*_*γ*_ is set to 0.5.

Fourth, inter-stimulus-intervals are independently sampled from a uniform distribution

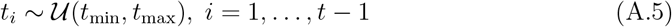

where *t*_min_ *∈ {*0, 2, 4*}* and *t*_max_ = *t*_min_ + 4.

Based on the sampled inter-stimulus-intervals (ISIs), the canonical hemodynamic response function (cHRF) as well as stimulus duration *t*_dur_ and repetition time TR, an *n × t* trial-wise design matrix *X*_*t*_ is generated which instantiates the sampled ISIs (see Figure 1A as an example for *t*_isi_ *∼ 𝒰* (0, 4)).

Moreover, an *n × n* temporal correlation matrix *V* is generated with entries

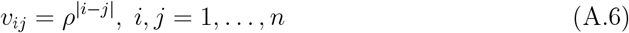

and a *v × v* spatial correlation matrix Σ is generated with entries

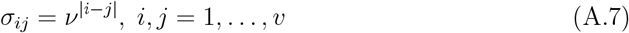

where the time constant is *ρ* = 0.12 and the space constant is *ν* = 0.48. This induced mild temporal correlations between subsequent scans *i* = 1, …, *n* and stronger spatial correlations between adjacent voxels *j* = 1, …, *v*.

Finally, an *n × v* noise matrix with standard deviation *σ ∈ {* 0.8, 1.6, 3.2*}* is sampled from the matrix-normal distribution with covariance matrices *V* and Σ

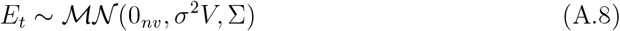

and the simulated fMRI signals are generated according to the trial-wise GLM as

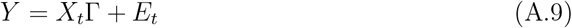

where the *t × v* matrix of trial-wise response amplitudes is given by

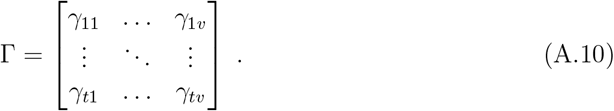

The combination of the 3 different options for *t*_min_/*t*_max_ and the 3 different options for *σ*^2^ leads to 9 different simulation scenarios (see Figures 3). In each scenario, *N* = 1,000 simulations with *S* = 2 sessions per simulation were performed.

After data generation, trial-wise activations are estimated and trial types are decoded. In the “least squares, all” (LS-A) approach, 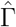 was obtained via equation (3) and a support vector classification was trained on the estimates from one session to predict trial types in the other session and vice versa. Decoding accuracy was quantified as the proportion of trials correctly assigned to trial types 1 and 2 in the test session. In the “least squares, separate” (LS-S) approach, 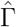 was obtained via equation (4) and the same support vector classification was applied. For inverse transformed encoding modelling (ITEM), 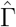 was obtained via equation (3) and trial types were decoded via ITEM-style inversion, with decoding accuracy being assessed as described in Section 2.5.

The present simulation differs from the one in Soch et al., 2020 in the following respects:

- The number of trials *t* was changed from 60 to 100 in order to increase the statistical power for all approaches compared to let *t* be sufficiently large in comparison with the number of voxels *v* = 33.
- The number of voxels was changed from 1 (univariate signal) to 33 (multivariate
- signals). This required to specify the spatial covariance for which we assumed that voxels come from a one-dimensional array and voxel noise is more correlated the closer the voxels are to each other (cf. eq. A.7).
- We replaced logistic regression (LR) by support vector classification (SVC) as the decoding algorithm, since LR becomes unstable for larger number of features and since SVC is more widely applied in neuroimaging data analysis.

### B A note on parametric modulators in SPM

The fMRI data set analyzed here represents a very unusual experiment in the sense that it can be understood as a single experimental condition (continuous visual stimulation) that is parametrically modulated with 48 variables (sector intensity levels) – which poses challenges for fMRI modelling.

When we started to analyze the data set with SPM, we noted that (i) parametric regressors were correlated to the onset regressor and (ii) parametric regressors are more correlated to each other than modulator variables from which they were generated (see Figure 7A). To us, both seemed to be unintended consequences, since onset and parametric regressors are typically aimed to model orthogonal components of the fMRI signal (condition mean vs. modulator effect) and parametric regressors should usually preserve the correlation of modulator variables.

We found out that this happens, because SPM does not transform modulator variables before HRF convolution, e.g. via mean-centering. Instead of mean-centering, SPM offers to orthogonalize regressors. Upon choosing this option, we noted that (i) parametric regressors were not correlated anymore, neither with the onset regressor nor among each other, but (ii) the closer one gets to the end of the design matrix (e.g. 48th vs. 1st parametric regressor), the lower is the correlation of parametric regressors with the original modulator variable (see Figure 7B).

We found out that this happens, because SPM uses sequential orthogonalization, i.e. the first parametric regressor is orthogonalized with respect to the onset regressors, the second parametric regressor is orthogonalized with respect to the first one, etc. (Ashburner et al., 2021). This has the consequence that, for a large number of parametric modulators, later parametric regressors are less and less veridical and interpretable. Specifically, only contrasts addressing all parametric regressors together are permitted, but not contrasts only looking at a subset of modulator variables.

We then went on to write our own function for creating an HRF-convolved design matrix from onsets, durations and parametric modulators^18^. Instead of sequential orthogonalization, we used mean-centering of each modulator variable to achieve orthogonality with the onset regressor. We found that our approach avoids both problems, i.e. (i) the correlation structure of the parametric regressors is close to that of the underlying modulator variables and (ii) interpretability of parametric regressors relative to modulator variables is preserved (see Figure 7D). The same was observed when mean-centering modulator variables before entering them into SPM and switching of SPM’s orthogonalization procedure which was revalidating our approach (see Figure 7C).

In sum, we therefore suggest – especially in designs with multiple modulator variables for an experimental condition – to turn off orthogonalization in SPM and to subtract the mean (or a neutral value, see below) from modulator variables before SPM model specification. This comes at the cost that parametric regressors may still be correlated, but it is worth paying the price, because this yields better interpretability of the parameter estimates for the parametric regressors (Mumford et al., 2015). Also, partial collinearity of parametric regressors – if it is due to the design – is valuable information that can be handled by linear model estimation (Vanhove, 2020).

In the case that the means of modulator variables are different across subjects (or between modulators), another option is to subtract a neutral value rather than the variable mean. For example, if stimulus ratings are collected from subjects during or after the experiment, those ratings will rarely be distributed uniformly for each subject (and likely be distributed differently between subjects). Then, it makes sense to subtract the value corresponding to the neutral rating from the modulator variable. In this case, it is possible that parametric regressors are correlated to their onset regressor, but in exchange, inter-subject interpretability is preserved. In the past, we have successfully applied this strategy to an fMRI episodic memory task (Soch et al., 2021b; Soch et al., 2021a) in which stimulus presentations were parametrically modulated with subsequent memory response ranging between 1 (“the stimulus is new”) and 5 (“the stimulus is old”), such that 3 (“I don’t know”) corresponded to the subtracted neutral response.

## Footnotes

1 Note that we use the word “scans” to refer to consecutive images acquired within one fMRI run (also called “TRs”), rather than to separate fMRI recording sessions (called “runs” or “sessions” [in SPM]).

2 See: https://statproofbook.github.io/P/tglm-dist.

3 See: https://statproofbook.github.io/P/iglm-dist.

4 Note that equation (9) is a multivariate GLM which entails two key assumptions: (i) linearity of the effects and (ii) normality of the errors. For a discussion of these assumptions, see Section 5.3.

5 Note that this step is not strictly necessary, as ITEM is based on a trial-wise, not condition-based GLM. However, the ITEM toolbox for SPM builds the trial-wise GLM by extracting trials from conditions encoded in the standard GLM, such that this step is recommended when working with SPM.

6 See function `ITEM_est_1st_level` in the repository https://github.com/JoramSoch/ITEM.

7 See functions `ITEM_dec_class` and `ITEM_dec_recon` of the ITEM toolbox.

8 See functions `ITEM_dec_class_SL` and `ITEM_dec_recon_SL` of the ITEM toolbox.

9 See function `GLMestimatesingletrial` in the repository https://github.com/cvnlab/GLMsingle.

10 See “Tips on Usage” in the GLMsingle wiki at https://glmsingle.readthedocs.io/en/latest/wiki.html#my-experiment-design-is-not-quite-synchronized-with-my-fmri-data.

11 See the third entry at https://oeis.org/A000605.

12 Also see: https://twitter.com/JoramSoch/status/1631568277800378368.

13 See Allefeld and Haynes, 2014, pp. 352-354 for a comprehensive discussion of these aspects.

14 URL: https://github.com/JoramSoch/ITEM.

15 URL: https://github.com/JoramSoch/ITEM-SL-paper.

16 URL: https://openneuro.org/datasets/ds002013.

17 See: https://github.com/JoramSoch/ITEM-SL-paper/blob/main/README.md.

18 See function `ITEM_get_des_mat` in the repository https://github.com/JoramSoch/ITEM.

